# A cranial implant for stabilizing whole-cell patch-clamp recordings in behaving rodents

**DOI:** 10.1101/2023.02.21.529357

**Authors:** Joshua Dacre, Michelle Sanchez-Rivera, Julia Schiemann, Stephen Currie, Julian J. Ammer, Ian Duguid

**Affiliations:** Centre for Discovery Brain Sciences and Patrick Wild Centre, Edinburgh Medical School: Biomedical Sciences, University of Edinburgh, Edinburgh, EH8 9XD, UK; Simons Initiative for the Developing Brain, University of Edinburgh, Edinburgh, EH8 9XD, UK; Center for Integrative Physiology and Molecular Medicine, Saarland University, Homburg, Germany; Optophysiology, University of Freiburg, Faculty of Biology, 79100 Freiburg, Germany

## Abstract

**Background:** *In vivo* patch-clamp recording techniques provide access to the sub- and suprathreshold membrane potential dynamics of individual neurons during behavior. However, maintaining recording stability throughout behavior is a significant challenge, and while methods for head restraint are commonly used to enhance stability, behaviorally related brain movement relative to the skull can severely impact the success rate and duration of whole-cell patch-clamp recordings.

**New method:** We developed a low-cost, biocompatible, and 3D-printable cranial implant capable of locally stabilizing brain movement, while permitting equivalent access to the brain when compared to a conventional craniotomy.

**Results:** Experiments in head-restrained behaving mice demonstrate that the cranial implant can reliably reduce the amplitude and speed of brain displacements, significantly improving the success rate of recordings across repeated bouts of motor behavior.

**Comparison with existing methods:** Our solution offers an improvement on currently available strategies for brain stabilization. Due to its small size, the implant can be retrofitted to most in vivo electrophysiology recording setups, providing a low cost, easily implementable solution for increasing intracellular recording stability in vivo.

**Conclusions:** By facilitating stable whole-cell patch-clamp recordings in vivo, biocompatible 3D printed implants should accelerate the investigation of single neuron computations underlying behavior.

## Introduction

Whole-cell patch-clamp recordings provide access to the membrane potential dynamics of individual neurons, and thus unique insight into the biophysical mechanisms that regulate input-output transformations. Although initially developed for in vitro applications (Neher and Sakmann 1976; Sigworth and Neher 1980; Hamill et al. 1981), the desire to understand how individual neurons process behaviorally-relevant information motivated the development of whole-cell patch-clamp recordings in anaesthetized (Margrie, Brecht, and Sakmann 2002; Pei et al. 1991), awake head-restrained (Margrie, Brecht, and Sakmann 2002; Harvey et al. 2009; Petersen et al., 2003) and even freely moving rodents (Lee et al. 2006; Lee, Epsztein, and Brecht 2009; Long and Lee 2012). Successful whole-cell recordings require the formation and maintenance of a high resistance seal between the glass electrode and the membrane of a neuron, which can be highly challenging in vivo. A common strategy to improve recording stability is to immobilize the rodent’s head with a lightweight headplate surgically implanted to the skull (Ono et al. 1985). However, movement of the brain relative to the skull, induced by either cardiac and respiratory pulsations (Dichter 1973; Avezaat and van Eijndhoven 1986; Paukert and Bergles 2012; Laffray et al. 2011) or body movement (Dombeck et al. 2007; Chen et al. 2013; Greenberg and Kerr 2009) can lead to irreparable loss of recording integrity (Kimura et al. 2012) (Figure 1).

**Figure 1.**
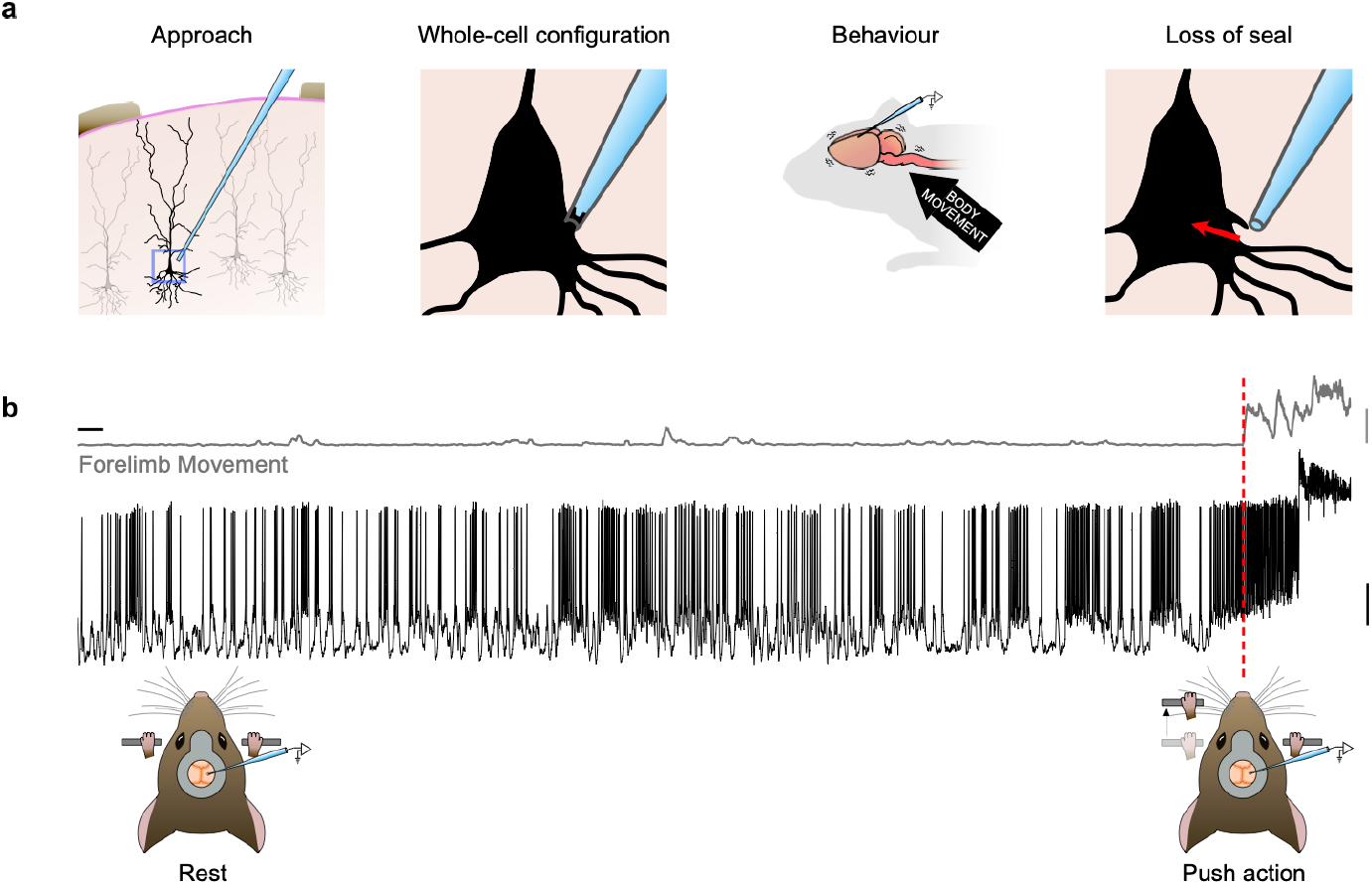
Movement-related loss of whole-cell recordings in vivo. (a) Left to right: schematic diagrams depicting approach of patch pipette, whole cell recording configuration and movement-related loss of seal integrity. (b) Top, example motion index trace describing gross movement of the mouse forelimb. Middle, membrane potential recording of a layer 5B pyramidal neuron in the caudal forelimb area of primary motor cortex. Bottom, schematic diagrams depicting forelimb movement during a cued forelimb lever push task. Red dashed line, movement initiation. Vertical scale bars, 1 arbitrary unit (grey, AU) and 20 mV (black), horizontal scale bar, 2s.

Several imaging-based approaches have been developed to account for, and minimize the impact of, brain movement using real-time movement-corrected 3D two-photon imaging (Griffiths et al. 2020), or trajectory adjustment of the approaching patch-pipette as a function of the motion of the target neuron (Suk et al. 2017; Annecchino et al. 2017). Such approaches involve mechanically and computationally intensive strategies, precluding their applicability to many experimental setups, including those lacking two-photon microscopes. The cost and complexity of two-photon-based solutions provides an additional barrier to exploring their potential use for stabilizing patchclamp recordings in behaving rodents.

Here we developed a 3D-printable cranial implant for stabilizing intracellular recordings in vivo, designed to meet the following criteria: the device must be simple to fabricate, made from biocompatible material and easy to surgically implant; it must improve the probability and longevity of patch-clamp recordings during behavior; and permit unhindered access to the brain region below to facilitate rapid topical application of pharmacological agents (Duguid et al., 2015; Schiemann et al., 2015). The surgical requirements are straightforward and similar to implanting a glass cranial window for in vivo 2-photon population imaging(Andermann et al., 2011; Dombeck et al., 2007; Holtmaat et al., 2009; Mostany and Portera-Cailliau, 2008; Roome and Kuhn, 2014), albeit much faster. By measuring medial-lateral and rostro-caudal brain displacements using two-photon imaging, we demonstrate that both the speed and amplitude of brain displacements were significantly reduced, increasing the success rate and longevity of patch-clamp recordings during repeated bouts of behavior. Due to its small size, low-cost, easily modifiable design and biocompatibility, our implant can be retrofitted to any rodent preparation and can be maintained for weeks to months.

## Methods

### Cranial implant design and manufacture

The 3-dimensional model of the cranial implant was designed in Fusion 360 (Autodesk) and fabricated from a biosafe plastic (Somos® BioClear or Somos® WaterShed XC 11122) using a stereolithographic 3-D printer (Viper SI2 SLA Printer) with a layer height of 100 μm. The main features of the implant are: a Ø 2.8 mm core designed to fit within a ~Ø 3 mm craniotomy; a core depth of 0.6 mm designed to exert light pressure on the surface of the brain; a Ø 3.6 mm rim to facilitate accurate placement and fixation to the skull; and a beveled well to allow recording pipettes access to the underlying cortex (Figure 2a). The diameter of the central hole (Ø 0.6 mm) was designed to accommodate most patch-clamp glass electrode geometries, even when patching from deep cortical layers or subcortical structures and is positioned off center so that when rotated the hole can be placed in a position which avoids any large underlying blood vessels (Figure 2b-c). The 3D drawings and dimensions of the implant can be accessed via https://doi.org/10.5281/zenodo.7632594.

**Figure 2.**
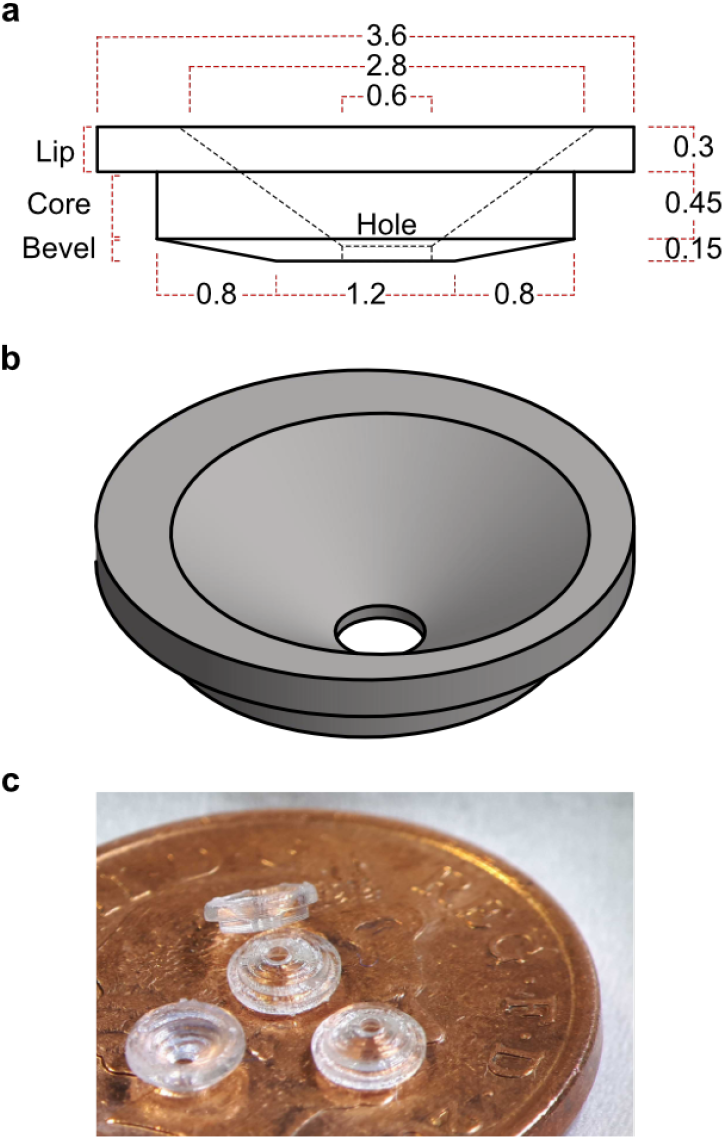
Implant design. (a) Dimensioned side- and (b) isometric view of the implant design. Measurements are in mm. (c) Four 3D printed implants placed on a UK one pence piece.

### Animals

Experiments were performed on adult male and female C57BL/6J wild-type (RRID: IMSR_ JAX:000664), and VGAT-Venus (Wang et al., 2009) mice (6 - 12 weeks old, 20-30 g, one to six animals per cage), maintained on a reversed 12:12 hour light:dark cycle (lights off at 7:00 am) and provided with ad libitum access to food and water. All experiments and procedures were approved by the University of Edinburgh local ethical review committee and performed under license from the UK Home Office in accordance with the Animal (Scientific Procedures) Act 1986.

### General Surgery

Mice undergoing surgery were induced with 4% and maintained under ~1.5% isoflurane anesthesia, with each animal receiving fluid replacement therapy (0.5 ml sterile Ringer’s solution) and buprenorphine (0.5 mg/kg; for pain relief), and the eyes covered by ointment (Bepanthen) to prevent drying. Additionally, buprenorphine (0.5 mg/kg) was administered in the form of an edible jelly cube within 24 hours of recovery from surgery. For surgeries involving removal of the dura, each animal received an injection of carprofen on the day of removal / recording (5 mg/kg). A small lightweight headplate (0.75 g) was implanted on the surface of the skull using cyanoacrylate super glue and dental cement (Lang Dental, USA), and the surface of the skull within the well of the headplate was covered with a thin layer of cyanoacrylate super glue to preserve bone health. Mice were then left for at least 48 hours to recover.

### Surgical implantation

On the day of recording, mice were anaesthetized and the headplate used to immobilize the head in a stereotaxic frame. Carprofen (5 mg/ kg) was administered at the start of surgery, and the eyes covered by ointment. The inside of the headplate was gently cleared with an air-duster and sterilized by gentle abrasion with a cotton bud soaked in ethanol. A handheld dentist drill (Ø 0.6 mm burr) was used to remove any adhesive material covering the skull above the region of interest, exposing an area of cleared skull >4 mm in diameter. A circular 3 mm diameter glass coverslip was placed on the skull, centered above the region of interest (caudal forelimb area (CFA): 1.6 mm lateral, 0.6 mm rostral to bregma), and the outline was scored into the skull using a 26-gauge needle attached to a 1 ml syringe (Figure 3a). Once the coverslip was removed, a dental drill was used to etch the outline of the craniotomy. The cranial implant was then placed onto the outline and any necessary alterations to the shape were made before thinning the skull along the etched outline (Figure 3b). Continuous movement of the drill head and intermittent irrigation with saline was essential to avoid unnecessary heating of the skull and to reduce the probability of underlying blood vessel rupture. Once thinned, the central bone area was removed with forceps (Dumont #5 forceps) avoiding disruption or removal of the underlying dura (Figure 3c). A 30-gauge needle attached to 1 ml syringe was used to perforate the dura and create a ~1 mm rostro-caudal slit in the center of the craniotomy, before fine forceps (Dumont 5SF forceps) were used to create a small, roughly circular durotomy (~1 mm in diameter) (Figure 3d).

**Figure 3.**
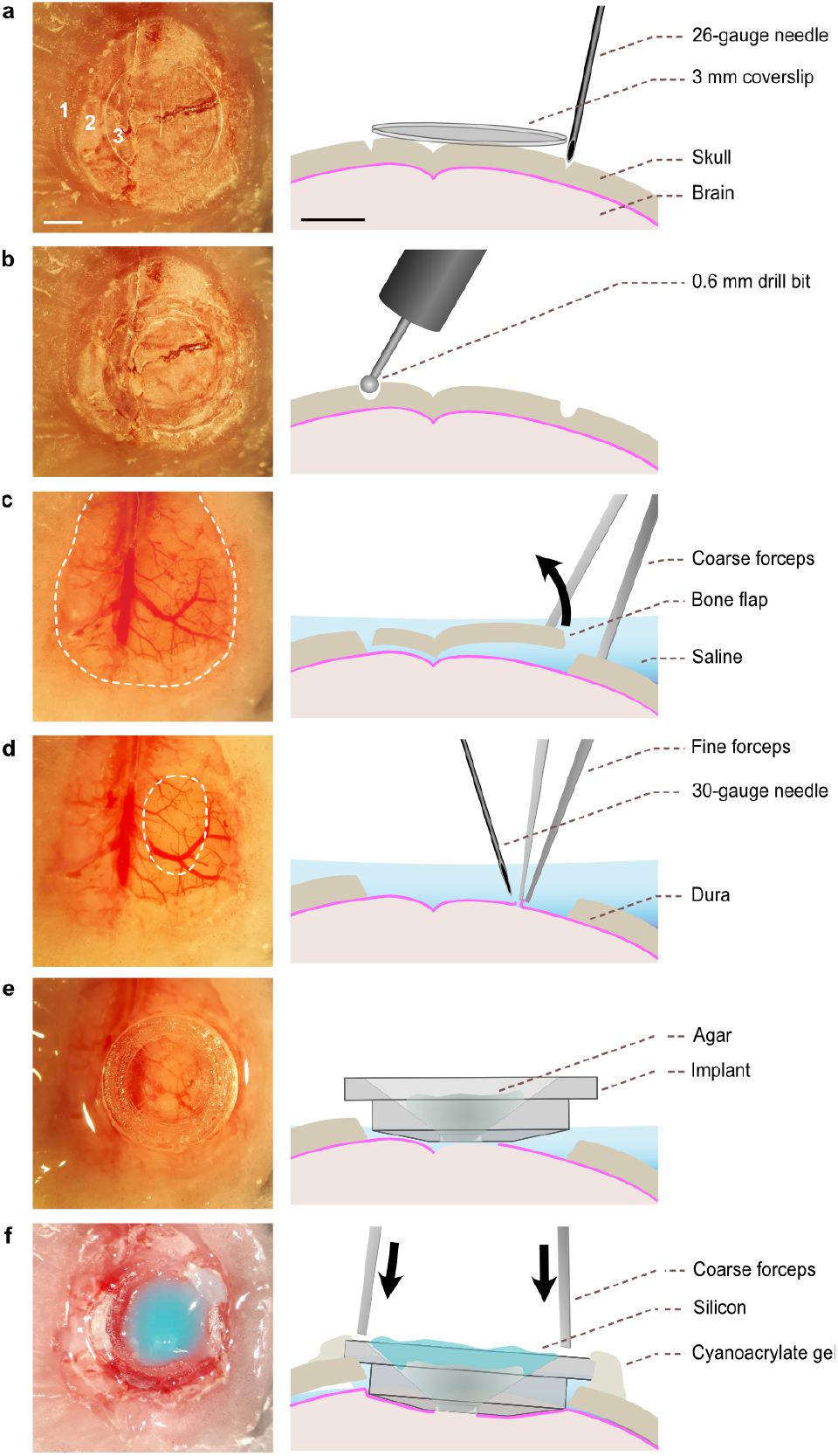
Surgical implantation. (a) Left, Image depicting the outline of a 3 mm glass coverslip, centred above the region of interest. Right, etching the circumference using a 26-gauge needle. 1 – dental cement affixing headplate to skull; 2 – exposed skull; 3 –glass coverslip. Scale bars, 1 mm. (b) Left, Image of the craniotomy outline. Right, a handheld drill with ⊘ 0.6 mm burr used to generate the craniotomy. (c) Left, Image of the craniotomy with central bone area removed (dashed area). Right, fine forceps used to remove bone while maintaining constant irrigation with saline. (d) Left, Image of the durotomy (dashed area). Right, 30-gauge needle and fine forceps are used to remove dura above targeted region. (e) Left, Placement of the implant. Right, implant is aligned to the craniotomy with central well filled with agar. (f) Left, Silicon sealed cranial insert. Right, Coarse forceps are used to press down and align the implant with the surface of the skull. After drying the remaining saline, gel superglue is used to affix the implant in place and silicon is added to the central well.

After irrigating any small vessel bleeds resulting from the durotomy (this step is not routinely required), the cranial implant was placed into the well, and most of the external solution removed while leaving enough to keep the exposed brain moist (Figure 3e). A small amount of agar was used to fill the well of the implant, with a thin layer of two-part silicone sealant added to prevent the agar from drying out. Coarse forceps (Dumont #4 forceps) were used to apply pressure evenly at the two opposite edges of the implant until the entire rim rested on the skull, and the remaining external solution was removed using a twisted tissue corner. Maintaining pressure with the forceps, a 30-gauge needle was used to apply gel cyanoacrylate superglue around the rim, which was left to dry for 10 mins (Figure 3f). Finally, two-part silicone was used to fill the entire of the headplate well to keep it clear of dust and debris during recovery and the mouse was placed in a temperature-controlled cage to recover for at least 1 hour. No adverse health effects were observed after implantation surgery.

### Cued forelimb lever push task

We assessed the ability of the cranial implant to improve patch-clamp recording stability using a cued lever push task for mice (Dacre et al., 2021). This behavior involves rapid forelimb, whole body, orofacial and tongue movements bracketing many of the actions likely to induce brain displacements during behavior.

Mice were handled extensively before being head-restrained and habituated to a custom lever push behavioral setup and trained to perform cued (6 kHz auditory tone) lever push movements to obtain a ~5 μl water reward (Dacre et al., 2021). To increase task engagement mice were placed on a water control paradigm (1 ml/day) and weighed daily to ensure body weight remained above 85% of baseline. Mice were trained once per day for 30 mins, with a quasi-random inter-trial-interval (ITI) of 4-6 s followed by presentation of the auditory cue. Mice were trained to respond within a10s response window early in training, reduced to 2 s across sessions, and were deemed ‘expert’ after achieving >90 rewards per session on two consecutive days. Lever movements during the ITI would result in a resetting of the lever and commencement of a subsequent ITI.

### Two-photon imaging

Priorto imaging neurons in the caudal forelimb region of motor cortex, mice were deeply an-aesthetized and immobilized in a stereotaxic frame. Carprofen (5 mg/kg) was administered by subcutaneous injection and the eyes covered by ointment to prevent them from drying out. Mice expressing the yellow fluorescent protein Venus under the vesicular GABA transporter (VGAT) promoter (Wang et al., 2009) underwent either a conventional craniotomy with dura left intact or removed, or insertion of a cranial implant with dura removed (Figure 4a and b). Mice were allowed to recover in a temperature-controlled cage for at least an hour before being head-restrained under the 2-photon microscope, at which point silicon and agar were removed, and the headplate was filled with external solution (containing: 150 mM NaCl, 2.5 mM KCl, 10 mM HEPES, 1 mM CaCl2, and 1 mM MgCl2 (adjusted to pH 7.3 with NaOH)). 2-photon imaging of Venus-expressing neurons was performed at three different cortical depth ranges (100-140μm, 300-340μm, 450-520μm below the pial surface) using a custom-built resonant scanning 2-photon microscope (320 x 320 μm FOV; 600 x 600 pixels) at 40 Hz frame rate, using a Ti:Sapphire pulsed laser (Chameleon Vision-S, Coherent, CA, USA; < 70 fs pulse width, 80 MHz repetition rate) tuned to 1020 nm wavelength (Figure 4c). Images were acquired with a 40xobjective lens (0.8 NA; Nikon) and custom-programmed LabVIEW-based software (LOTOS scan).

**Figure 4.**
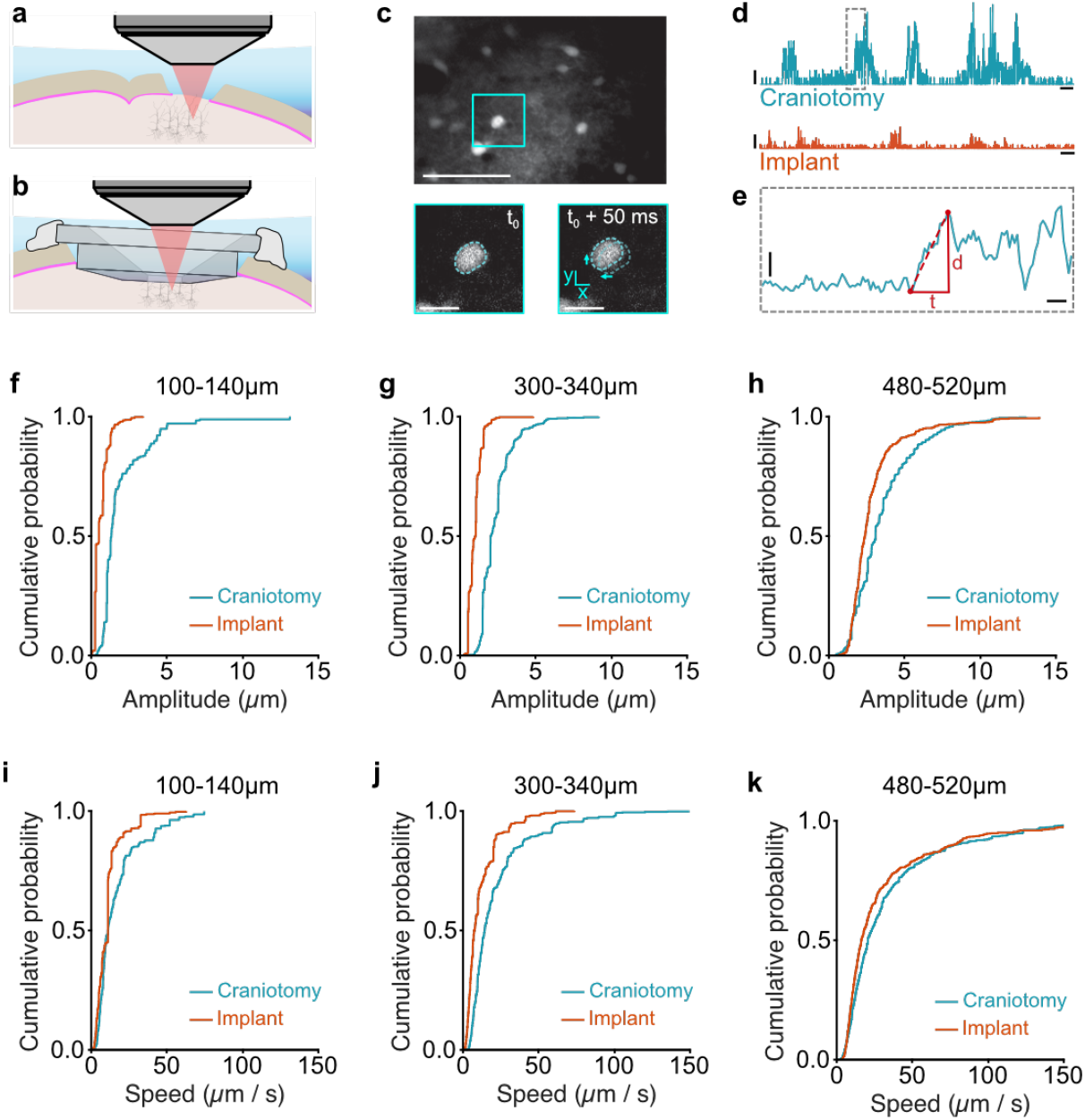
Implant reduces speed and amplitude of brain displacements. Two-photon imaging of L2/3 pyramidal neurons through (a) a conventional craniotomy or (b) an implant. (c) Top, Example 2-photon imaging field-of-view from layer 2/3 in CFA (average time projection of 1 s of raw data, peri-movement initiation; scale bar, 50 μm). Bottom, expanded view of cyan square showing example L2/3 neuron before and after (average time projection of 4 frames or 100 ms of raw data) the onset of movement. Note xy shift in location (cyan dotted lines). Bottom scale bars, 10 μm. (d) Example trace showing brain displacements from the point of origin across time in the absence (top trace) and presence (bottom trace) of an implant. Horizontal scale bars, 2 s; vertical scale bars, 2 μm. (e) Expanded view of grey dashed rectangle in (d) showing the time (t) and distance (d) of an individual displacement. Speed (s) = distance (d) / time (t). Horizontal scale bar, 200 ms; vertical scale bar, 1 μm. (f-h) Cumulative probability plots showing the distribution of brain displacement amplitudes using a conventional craniotomy with dura removed (teal, N = 3 mice) or implant (orange, N = 3 mice) at 3 different depths from the pial surface (100-140μm, 300-340μm, 480-520μm). (i-k) Cumulative probability plots showing the distribution of brain displacement speeds using a conventional craniotomy with dura removed (teal, N = 3 mice) or an implant (orange, N = 3 mice) at 3 different depths from the pial surface (100-140μm, 300-340μm, 480-520μm).

### Whole-cell patch-clamp electrophysiology

Priorto whole-cell patch-clamp recording, mice were deeply anaesthetized and immobilized in a stereotaxic frame. Carprofen (5 mg/kg) was administered by subcutaneous injection and the eyes covered by ointment. For comparison, mice underwent either a conventional craniotomy with dura removed or insertion of a cranial implant with dura removed. Glass patch pipettes were pulled to a resistance of 5.5 - 7.5 MΩ (number of pipettes per experiment: craniotomy 6.4 [5.2 7.4] 95% CI, N = 56 mice; implant 7.4 [6.3 8.4] 95% CI, N = 84 mice), filled with internal solution (285–295 mOsm, containing: 135 mM K-gluconate, 4 mM KCl, 10 mM HEPES, 10 mM sodium phosphocreatine, 2 mM MgATP, 2 mM Na2ATP, 0.5 mM Na2GTP, and 2 mg/ml biocytin, pH adjusted to 7.2 with KOH), were lowered to 600 μm below the pial surface (layer 5) at an angle of 30° from vertical, under 300 mBar positive pressure at ~30-50 μm/s using a micromanipulator (PatchStar, Scientifica), at which point positive pressure was reduced to 20 mBar ‘searching pressure’. Using a test pulse (−10 mV step) visualized on an oscilloscope, the pipette was advanced in discrete steps of 2 μm at a rate of approximately 1 step / sec. Discrete step-like decreases in test pulse amplitude occurred when in close apposition to a cell membrane termed a ‘hit’. After 3 hits, stepping ceased and 5-10 mBar of negative pressure and a holding potential of −20 mV was applied. To assist gigaseal formation the holding potential was gradually lowered to −60 mV at a rate of ~1-5 mV/s. Signals were acquired at 20 kHz using a Multiclamp 700B amplifier (Molecular Devices) and filtered at 10 kHz using PClamp 10 software in conjunction with a DigiData 1440 DAC interface (Molecular Devices). No biascurrentwas injectedduring recordingsand the membrane potential was not corrected for junction potential. After each recording the patch pipette was retracted to form an outside-out patch and mice were anaethestised, transcardially perfused with paraformaldehyde, sliced, mounted, and imaged to recover and identify the location of each cell.

## Results

### Reduced brain displacement

To assess how the cranial implant affected brain displacements, we imaged Venus-expressing GABAergic interneurons in CFA through a conventional craniotomy in which the dura was left intact (dura on, N = 3 mice), the dura was removed (dura off, N = 3 mice) or through a cranial implant with the dura removed (cranial implant, N = 3 mice) (Figure 4a-c). We assessed brain displacements at three different cortical depths covering L1, L2-4 and L5 (100-140 μm, 300-340 μm, 450-520 μm from the pial surface). Motion artefacts in the raw fluorescence timeseries corresponding to brain displacements were measured with the rigid motion correction capabilities of Sequential Image Analysis (SIMA), an Open Source package for the analysis of time-series imaging data (Kaifosh et al., 2014). Custom written MATLAB scripts were used to analyze the frame-by-frame x,y-displacement vectors calculated by SIMA by converting them to a scalar distance from the origin using Pythagoras’ theorem (Figure 4d-e). Threshold crossings, defined as the 10th percentile of all displacement values acquired during a single recording, were used to categorize episodes of movement versus non-movement and individual movement periods were considered separate if they occurred with an inter-event interval of more than 500 ms. Movement episodes tended to consist of a rapid initial displacement, followed by a gradual return to the starting location (Figure 4d-e). Given that rapid brain displacements are likely to cause loss of seal integrity, we focused our analysis on the initial displacement amplitude and speed. The amplitude was defined as the largest displacement in a 500 ms window around the point of threshold crossing, while the duration was defined as the time from baseline to peak (Figure 4e).

Brain displacements measured using a conventional craniotomy with dura removed were between 1-15 μm across all cortical depths. By applying a cranial implant, the speed, amplitude, and duration ofdisplacements were significantly reduced. Stabilization was highest in superficial layers and reduced as a function of distance from the pial surface (100-140 μm = 68%, 300-340 μm = 59%, 480-520 μm = 19% reduction in displacement amplitude) (Table 1, Figure 4f-k). This suggests that the implant provides a stabilization point on the surface of the brain, that is effective in reducing brain displacements across cortical layers. Interestingly, imaging through a conventional craniotomy with the dura intact, similarto using collagenase to thin the dura prior to recording (Zhu et al., 2002), actually increased the speed and amplitude of displacements when compared to the same preparation but with dura removed (Table 1).The increased stability after performing a durotomy likely relates to the reequilibration ofintracranial pressure which stabilizes the brain within the bone removal area. Adding a cranial implant then further reduces the speed and amplitude of brain displacements to further stabilize the brain.

**Table 1:** Speed, amplitude, and duration of movement-related brain displacements.

| Configuration | (n) | Amplitude ($\mu\text{m}$) | Speed ($\mu\text{m/s}$) | Duration (s) |
| --- | --- | --- | --- | --- |
| **[100-140 $\mu\text{m}$]** |  |  |  |  |
| Dura on | $n = 204$ | 2.7 [1.8 3.7] | 20.0 [14.8 25.7] | 0.17 [0.14 0.20] |
| Dura off | $n = 309$ | 2.0 [1.6 2.4] | 16.3 [12.8 20.3] | 0.18 [0.15 0.22] |
| Cranial implant | $n = 526$ | 0.6 [0.5 0.7] | 10.3 [8.8 12.0] | 0.10 [0.08 0.12] |
| % reduction | (off vs implant) | 68.4% | 36.7% | 44.4% |
| KS test ($p$ value) | (off vs implant) | **$1.56 \times 10^{-87}$** | **$7.02 \times 10^{-15}$** | **$7.94 \times 10^{-16}$** |
| **[300-340 $\mu\text{m}$]** |  |  |  |  |
| Dura on | $n = 402$ | 3.4 [2.6 4.4] | 31.1 [23.1 41.4] | 0.16 [0.14 0.19] |
| Dura off | $n = 488$ | 2.4 [2.2 2.6] | 23.0 [18.8 27.8] | 0.17 [0.15 0.19] |
| Cranial implant | $n = 462$ | 1.0 [0.9 1.0] | 12.2 [10.7 13.9] | 0.13 [0.12 0.15] |
| % reduction | (off vs implant) | 58.9% | 47.1% | 25.5% |
| KS test ($p$ value) | (off vs implant) | **$2.20 \times 10^{-189}$** | **$4.94 \times 10^{-32}$** | **$2.26 \times 10^{-5}$** |
| **[480-520 $\mu\text{m}$]** |  |  |  |  |
| Dura on | $n = 352$ | 4.7 [4.2 5.2] | 49.9 [41.6 58.9] | 0.15 [0.14 0.17] |
| Dura off | $n = 452$ | 3. [3.22 3.8] | 35.2 [29.6 41.7] | 0.17 [0.16 0.19] |
| Cranial implant | $n = 399$ | 2.86 [2.6 3.2] | 31.8 [25.6 38.7] | 0.16 [0.15 0.18] |
| % reduction | (off vs implant) | 18.8% | 9.9% | 5.9% |
| KS test ($p$ value) | (off vs implant) | **$7.70 \times 10^{-51}$** | **$6.04 \times 10^{-15}$** | 0.186 |

### Improved whole-cell recording stability during behavior

Our imaging data demonstrated that the cranial implant suppresses brain displacements across cortical layers, but the magnitude is depth dependent with deeper layers showing less suppression. To investigate whether this ‘dampening’ effect is translated into increased success rate and longevity of recordings, we performed blind patch-clamp recordings from layer 5B pyramidal neurons, either through a conventional craniotomy or a cranial implant in mice trained to perform our lever push task (Figure 5a-b). The implant did not affect the probability of forming a gigaseal (P(Gigaseal): craniotomy 0.27 [0.17 0.40] 95% CI, N = 56 mice; implant 0.31 [0.23 0.40] 95% CI, N = 84 mice), or attaining a whole-cell recording configuration (P(break-in): craniotomy 0.62 [0.41 0.82] 95% CI, N = 56 mice; implant 0.61 [0.47 0.74] 95% CI, N = 84 mice). In a subset of neurons we recorded the mean resting membrane potential immediately after break-in and found it was not different across preparations (Vrest: craniotomy −60.7 mV [−68.2 −49.2] 95% CI, n = 28 neurons, N = 28 mice; implant −62.3 mV [−66.5 −58.2] 95% CI, n = 59 neurons, N = 59 mice, Mann Whitney test p = 0.96) (Figure 5c), suggesting neuronal health was not adversely affected, and on average series resistances were consistently lower (Rs: craniotomy 49.2 MΩ [40.8 59.3] 95% CI, n = 28 neurons, N = 28 mice; implant 41.6 MΩ [36.6 48.7] 95% CI, n = 59 neurons, N = 59 mice, Mann Whitney test p = 0.003) (Figure 5d).

**Figure 5.**
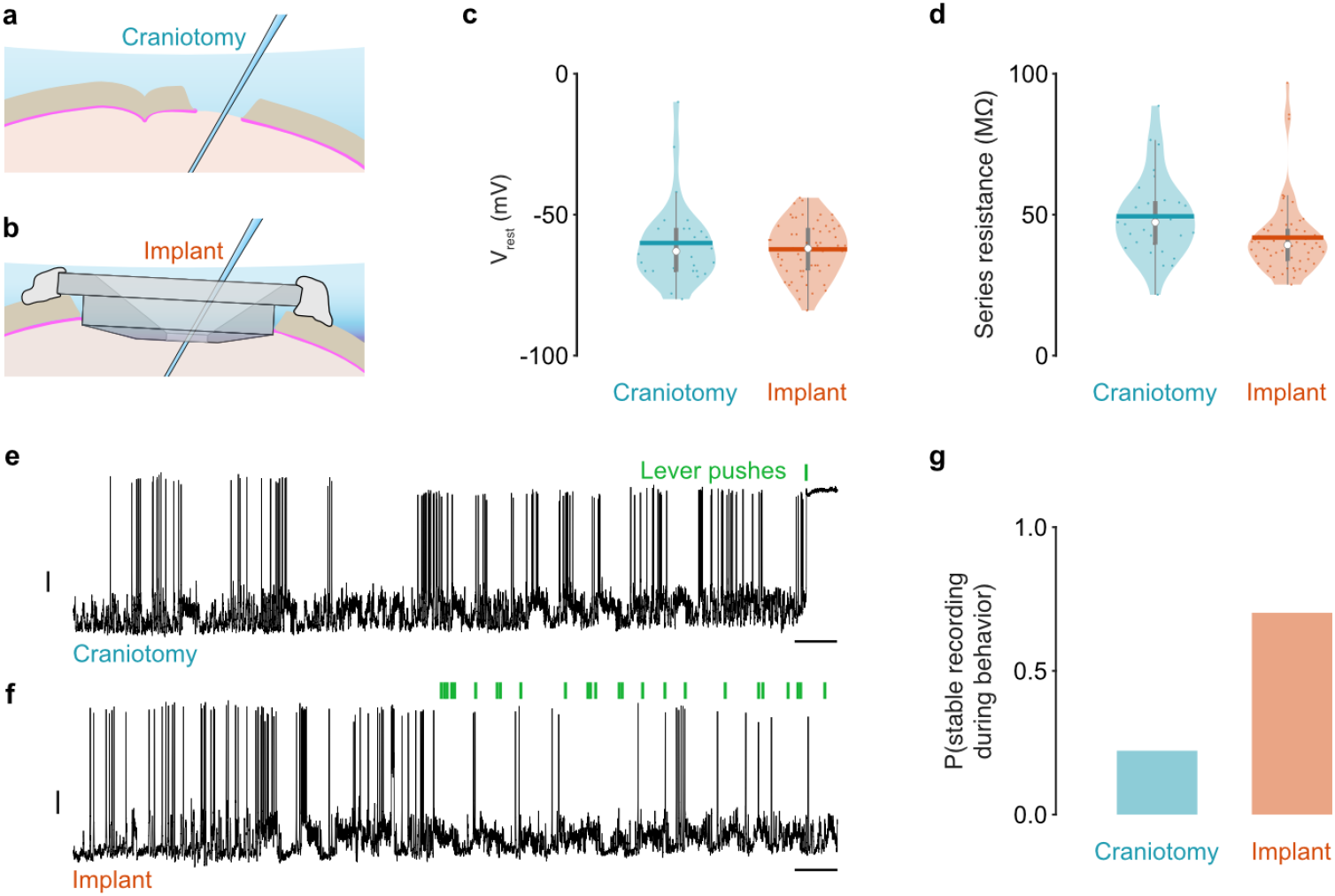
Cranial implant improves whole-cell recording stability. (a) Schematic showing patch-clamp recording through a conventional craniotomy. (b) Schematic showing patch-clamp recording through an implant. (c) Violin plots showing median (white circle), mean (thick horizontal line), inter quartile range (thick grey vertical line) and range (thin grey vertical line) of resting membrane potentials (Vrest) using a conventional craniotomy or implant. Dots represent data from individual mice (N = 28 and 59 mice, respectively). (d) Violin plots showing median (white circle), mean (thick horizontal line), inter quartile range (thick grey vertical line) and range (thin grey vertical line) of series resistance measured during whole cell recording using a conventional craniotomy or implant. Dots represent individual recordings from individual mice (N = 28 and 59 mice, respectively). (e) Membrane potential recording from a layer 5B pyramidal neuron using a conventional craniotomy. Green vertical bars represent lever pushes. Black vertical scale bar, 10 mV; horizontal scale bar, 5 s. (f) Membrane potential recording from a layer 5B pyramidal neuron using a cranial implant. Green vertical bars represent lever pushes. Black vertical scale bar, 10 mV; horizontal scale bar, 5 s. (g) Bar graph showing the probability of maintaining a stable whole-cell recording configuration during behavior when using a conventional craniotomy or implant.

To investigate whether the cranial implant increased the success rate and longevity of recordings in behaving mice, we recorded the membrane potential of a subset of layer 5B neurons that displayed stable membrane potential and spike heights at rest. We found that during behavior recordings using a conventional craniotomy were unstable and inevitably led to an abrupt loss of seal integrity upon movement initiation (n = 4/18, 22.2% of recordings were stable). In contrast, recording through the implant ensured whole-cell recordings were maintained throughout repeated bouts of behavior, which involved rapid forelimb, whole body, orofacial and tongue movements (n = 33/47, 70.2% of recordings were stable) (chi-squared test p = 0.0013) (Figure 5e-g). Importantly, this significant increase in recording stability likely represents a lower bound given that in layer 2/3 brain displacements were reduced by ~70% as opposed to ~30% in layer 5B. Although some cranial implant recordings were purposely terminated after only a few trials of behavior, in the remainder we compared the recording duration and average number of lever pushes in the absence and presence of the cranial implant. Stabilizing the brain resulted in a significant increase in both recording duration (craniotomy 5.3 mins [3.1 8.3] 95% CI, range [1 15] mins, N = 18 mice; implant 10.2 [7.2 14.1] 95% CI, range [3 28] mins, N = 33 mice, Mann Whitney test, p = 0.0005) and number of behavioral trials before seal integrity was lost (craniotomy 0.4 pushes [0 14.8] 95% CI, range [0 57] pushes, N = 18 mice; implant 34.1 [10.7 65.1] 95% CI, range [1 149] pushes, N = 33 mice, Mann Whitney test, p = 0.000001). This demonstrates a significant improvement permitting the stable recording of membrane potential dynamics across repeated bouts of motor behavior.

## Discussion

Whole-cell patch-clamp recordings remain the gold standard for investigating input-output transformations of individual neurons, but due to mechanical instability performing such recordings in vivo remains highly challenging. Here, we designed a cranial implant that is: simple to fabricate using 3-D printable biocompatible materials; straightforward to surgically implant; and permits unhindered access to the underlying brain region, which collectively improve the probability and longevity of recordings in behaving rodents. Allowing unhindered access to the underlying cortical surface permits the rapid topical application of pharmacological agents (Duguid et al., 2015; Schiemann et al., 2015), significantly reducing drug diffusion times when compared to application in the presence of agar or thinned dura (Jordan, 2021; Margrie et al., 2002; Zhu et al., 2002).

Reducing mechanical instability when performing electrophysiological recordings in vivo requires a two-step process. Firstly, stabilization of the head using custom milled (Dombeck et al., 2007; Jordan, 2021; Petersen, 2017) or 3-D printed head plates (Osborne and Dudman, 2014) minimizes lateral or axial displacement of the skull with respect to the recording electrode. Such an arrangement blocks head movements and translation of torsional forces generated during behavior which, if left unchecked, would significantly reduce the probability and duration of any recording. Although methods for whole-cell patch-clamp recordings in freely moving rodents do exist (Lee et al., 2009; Lee et al., 2006; Lee et al., 2014), they remain highly challenging and result in short duration recordings that may not be suitable for investigating the neural dynamics of many learned behaviors. Secondly, and perhaps less well described, is the minimization of brain movement with respect to the skull, providing an additional layer of stabilization necessary for maintaining recordings throughout behavior. Brain displacements arise due to cardiac and respiratory pulsations (Dichter 1973; Avezaat and van Ejndhoven 1986; Paukert and Bergles 2012; Laffray et al. 2011) and from torsional forces that propagate from the body, via the brainstem, to the brain. While several mechanically and computationally intensive strategies capable of accommodating brain movement currently exist (Suk et al. 2017; Annecchino et al. 2017; Griffiths et al. 2020), their cost and complexity create significant obstacles in terms of implementation in pre-existing experimental configurations. Moreover, simpler approaches to minimize brain movement using glass cranial windows (Andermann et al., 2011; Dombeck et al., 2007; Holtmaat et al., 2009; Mostany and Portera-Cailliau, 2008; Roone & Kuhn, 2014) provide only limited utility for patch-clamp electrophysiology. Our 3-D printable implant solution addresses the technical challenge of ensuring brain stability while also facilitating electrophysiological recordings in vivo. The implant was designed to apply gentle, prolonged downward pressure across a large surface are of the brain (2.8 mm), providing maximal tissue stability in the location where recordings will be performed. Given that the implant did not adversely affect the probability of attaining a whole-cell configuration, average series resistance or resting membrane potential of cortical neurons it is likely that the basic cytoarchitecture and function of the underlying tissue was preserved.

The simple design of our cranial implant enables it to be retrofitted to any pre-existing in vivo patch-clamp setup, without the need for restructuring or redesigning the head restraint apparatus. Unlike other approaches that use custom-milled glass coverslips (Andermann et al., 2011; Dombeck et al., 2007; Holtmaat et al., 2009; Mostany and Portera-Cailliau, 2008; Roone & Kuhn, 2014), access to the underlying brain structure is facilitated by an off-centered hole which is sufficiently wide to allow most patch pipette geometries to pass unhindered, while small enough not to compromise stability of the underlying tissue. By placing the hole off-center, the experimenter has the option to rotate the implant, and thus the pipette entry point, to avoid any underlying blood vessels. This is an important consideration given that blood vessel rupture is one of the main causes of pipette blockages and reduced gigaseal formation when performing whole-cell patchclamp recordings in vivo (Jordan, 2021; Petersen, 2017). To enable reproduction, and customization of the implant to individual applications, the design has been included in supplementary materials, and while many stereolithographic 3D printing materials are currently available, the biosafe plastics (Somos® BioClear or Somos® WaterShed XC 11122) used in this study will likely facilitate stable implantation for weeks to months depending on biological application.

While other influences certainly contribute to recording stability (e.g., skill of the experimenter, pipette geometry, body posture, age, and behavioral state of the animal), the fact that modest reductions in horizontal brain displacements after implantation led to such striking improvements in recording success suggests that it is medium to large amplitude, rapid brain movements that lead to irreparable loss of seal integrity. Although we only investigated stability in the context of a single behavior, our cued forelimb push task involved whole body repositioning, rapid limb extension / retraction, whisking, and orofacial / tongue movements (Dacre et al., 2021). As such, we expect a similar range of brain displacement kinematics across other head restrained rodent behaviors (Dombeck et al., 2007; Galinanes et al., 2018; Guo et al., 2015; Kislin et al., 2014; Peters et al., 2014; Schiemann et al., 2015) and anticipate the cranial implant to offer similar benefits regardless of experimental application or electrophysiological recording method.

## Author Contributions

Conceptualization and design, J.D., M.S.R,J.S., and I.D.; methodology & investigation, J.D., S.C., M.S.R., and J.J.A.; resources, S.C., M.S.R., and J.J.A.; writing – review & editing, all authors.

## Data Availability

The code for imaging analysis, including SIMA motion correction, can be found at https://git-hub.com/losonczylab/sima. Code and data used to generate paper figures will be provided upon reasonable request. The code is implemented in Matlab (2021b, MathWorks).

## Acknowledgment

We are grateful to members of the Duguid lab for experimental discussions and comments on the manuscript. VGAT-Venus mice were a gift from Yuchio Yanagawa, Gunma University Graduate School of Medicine, Maebashi, Japan, and Venus was developed by Dr. Atsushi Miyawaki at RIKEN, Wako, Japan. This study was supported by the BBSRC (UK)(BB/ R018537/1), DFG (Germany) (SCHI1267/2-1 to J.S. and AM 443/1-1 to J.J.A.), the Shirley Foundation (UK), and a Wellcome Senior Research Fellowship (UK) (110131/Z/15/Z) to I. D.

